# Assembly of the Bacterial Ribosome with Circularly Permuted rRNA

**DOI:** 10.1101/2024.04.10.588894

**Authors:** Xiyu Dong, Kai Sheng, Luca F.R. Gebert, Sriram Aiyer, Ian J. MacRae, Dmitry Lyumkis, James R. Williamson

## Abstract

Co-transcriptional assembly is an integral feature of the formation of RNA-protein complexes that mediate translation. For ribosome synthesis, prior studies have indicated that the strict order of transcription of rRNA domains may not be obligatory during bacterial ribosome biogenesis, since a series of circularly permuted rRNAs are viable. In this work, we report the insights into assembly of the bacterial ribosome large subunit (LSU) based on cryo-EM density maps of intermediates that accumulate during *in vitro* ribosome synthesis using a set of circularly permuted (CiPer) rRNAs. The observed ensemble of twenty-three resolved ribosome large subunit intermediates reveals conserved assembly routes with an underlying hierarchy among cooperative assembly blocks. There are intricate interdependencies for the formation of key structural rRNA helices revealed from the circular permutation of rRNA. While the order of domain synthesis is not obligatory, the order of domain association does appear to proceed with a particular order, likely due to the strong evolutionary pressure on efficient ribosome synthesis. This work reinforces the robustness of the known assembly hierarchy of the bacterial large ribosomal subunit, and offers a coherent view of how efficient assembly of CiPer rRNAs can be understood in that context.

## INTRODUCTION

Ribosome biogenesis is fundamentally an RNA folding process, and the efficiency of ribosome formation is a key aspect of fitness and evolutionary pressure. The precise folding of rRNA, formation of RNA tertiary interactions, and binding with associated proteins are all essential for ribosomal subunit assembly and subsequent protein synthesis. In previous *in vitro* studies, the interaction between pioneer ribosomal proteins and specific 5’ domains of the 23S rRNA was found to initiate the ribosome 50S subunit assembly (1-3). This observation led to the hypothesis that the 50S assembly is linked with the transcription of 23S rRNA, progressing from the 5’ to 3’ end. Recent research has demonstrated that the earliest assembly of bacterial 23S rRNA can be traced back to an immature LSU precursor comprising 500-600 nucleotides of the 5’-end of 23S rRNA and three ribosomal proteins (4-6). In our laboratory, we also proposed a mechanistic framework where the 50S subunit assembles hierarchically from the 5’-end to the 3’-end (5), indicating a coordinated assembly process with rRNA transcription.

Four circular permutants of *E. coli* rRNA were identified that are capable of folding into the correct conformation and supporting protein synthesis in the *E. coli* strain Δ7 *prrn*, with only a slight increase in the growth doubling time (7). At that time, the details of the assembly process were limited to information from the Nierhaus map (1) and the structures of the 50S ribosomal subunit (8,9), and it was difficult to understand the mechanistic implications of this intriguing observation. In addition, a circular permutant of rRNA from *T*.*aquaticus* was employed to introduce chemical modifications into the PTC region (10). Natural instances of circular permutation in rRNA molecules include the 23S rRNA of *Pyrococcus furiosus* (11) and the LSU rRNA of the mitochondria in *Tetrahymena pyriformis* (12). Inspired by the successful function of these circularly permuted RNAs, the Ribo-T was engineered, which is a ribosome with tethered subunits for use in synthetic biology (13,14). Ribo-T features a 23S rRNA that was circularly permuted at Helix 101, and was continuously connected to two 16S rRNA fragments at 5’-end and 3’-end.

These studies and observations challenge our view of co-transcriptional assembly in the normal cases, and suggest that rRNA possesses remarkable flexibility in coping with major perturbations. Our laboratory’s preceding structural analysis proposed a modular and parallel assembly model for the 50S subunit, organized based on the hierarchical docking of cooperative blocks (5,15), where the earliest stages of assembly do parallel the order of domain transcription. Consequently, we sought to determine whether the rRNA circular permutants adhere to the same assembly hierarchy or strictly follow the 5’-3’ co-transcriptional folding. In these circular permutants, the transcription starting sites differ from those of the wild type rRNAs. If the 50S subunit strictly undergoes co-transcriptional assembly, alternative immature assembly intermediates are expected to be observed during the ribosome biogenesis process.

In this study, four circularly permuted (CiPer) rRNAs were adapted for use in integrated ribosome Synthesis, Assembly and Translation (iSAT) (16). To investigate their assembly mechanism further, we determined the cryo-EM structures of the ribosome intermediates composed of circularly permuted rRNA that accumulate during ribosome synthesis *in vitro*. The resolved LSU intermediates exhibited some specific features specific to rearranged rRNA sequences. However, in general, the circularly permuted rRNAs go through the same assembly hierarchy that was previously established for synthesis of native ribosomes *in vitro* and in cells (5,6,15). This work illuminates the further folding logic of the rRNA and confirms the robustness of the original biogenesis pathways of the bacterial ribosome, as a driver of efficient ribosome synthesis.

## MATERIALS AND METHODS

### Purification of S150 extract and total protein of 70S ribosomes (TP70)

Native ribosomes and S150 extract were purified from S30 crude cell extract as described (17). Briefly, *E. coli* MRE cells were lysed, and S30 crude extract was obtained by two clarification spins. The S30 crude extract was layered onto a 37.5% sucrose cushion and ultracentrifugation at 90,000 g and 4 °C for 18 h, resulting in a ribosome containing pellet and S150 containing supernatant. Ribosomes were resolved in buffer C (10 mM TrisOAc (pH 7.5 at 4 °C), 60 mM NH_4_Cl, 7.5 mM Mg(OAc)_2_, 0.5 mM EDTA, 2 mM DTT) and stored at −80 °C. The concentration of 70S ribosomes was determined by A_260_readings (1 A_260_ unit = 24 pmol 70S) (18).

The supernatant was centrifuged at 150,000 g and 4 °C for 3 h and the top two-thirds of the supernatant was recovered. Using Snake Skin Dialysis tubing (3500 Da MWCO, Thermo fisher) The S150 extract was dialyzed into the AEB storage buffer (10 mM TrisOAc, pH 7.5 at 4°C, 20 mM NH_4_OAc, 30 mM KOAc, 200 mM KGlu,10 mM Mg(OAc)_2_, 1 mM spermidine, 1 mM putrescine, 1 mM DTT) and concentrated using a 3 kDa MWCO Centriprep concentrators (Merck Millipore). For best performance in the iSAT reaction, the A_260_ and A_280_ of the S150 extract was targeted to be 25 OD and 15 OD, respectively. The S150 extract was stored in small aliquots at −80 °C.

Ribosomal proteins were purified using existing protocols (17). The concentration of TP 70 was determined by measuring A_230_ (1 A_230_ unit = 240 pmol TP70) by SmartSpec Plus spectrophotometer (BIORAD). Aliquots were flash frozen and stored at −80°C.

### iSAT reaction

The iSAT reaction was performed as described previously (16,17) with slight modifications on the protein concentration and the operon of plasmid. Briefly, the iSAT reaction was performed in 57 mM Hepes-KOH, 1.5 mM spermidine, 1 mM putrescine, 10 mM Mg(Glu)_2_, and 150 mM KGlu at pH 7.5 with 2 mM DTT, 0.33 mM NAD, 0.27 mM CoA, 4 mM sodium oxalate, 2% w/v PEG-6000, 2 mM amino acids (Sigma-Aldrich), 1 nM pY71sfGFP plasmid encoding superfolder GFP (M. Jewett), 0.1 mg/mL T7 RNA polymerase, 42 mM phosphoenolpyruvate (Roche) and NTP+ mix (1.6 mM ATP (Sigma), 1.15 mM of GTP, CTP and UTP each (Sigma), 45.3 μg/μL tRNA from *E. coli* MRE 600 (Roche), 227.5 μg/μL Folinic acid. The above components were premixed. A 5.36 µL aliquot of the premix was pipetted into 5 µl of S150 extract. Ribosomal proteins and plasmid encoding the engineered rrnB operon were added to a final concentration of 0.6 µM and 4 nM respectively. iSAT reactions of 15 μL each were performed in 96 well plates (Applied Biosystems) and incubated in a CFX Connect Real-time System (BIORAD) at 37 °C for variable time. sfGFP production was detected by fluorescence measurement at 5 min intervals (excitation: 450-490 nm, emission: 510-530 nm).

### Electron microscopy sample preparation

iSAT reactions were diluted approximately three-fold with buffer A (50 mM Tris pH 7.8, 10 mM MgCl_2_, 100 mM NH_4_Cl, 6 mM β-mercaptoethanol) and loaded onto a 10-40% (w/v) sucrose gradient in buffer E. Gradients were spun in a Beckman SW41 rotor at 26,000 rpm for 16 h at 4 °C. Gradient fractions containing LSU particles as indicated by A_260_ readings and agarose gel were spin-concentrated using a 100 kDa MW cutoff filter (Amicon) and the buffer was exchanged to buffer A (20 mM Tris-HCl, 100 mM NH_4_Cl, 10 mM MgCl_2_, 0.5 mM EDTA, 6 mM β-mercaptoethanol; pH 7.5). 3 µL sample was applied to a plasma cleaned (Gatan, Solarus) Quantifoil R 1.2/1.3 2 nm carbon coated copper grid and manually plunge frozen in liquid ethane (19).

### Electron microscopy data collection

Single-particle data were collected using Leginon software (20) on a Talos Arctica transmission electron microscope equipped with a K2 direct electron detector, operating at 200 keV with pixel size of 1.15 Å at 36,000 magnification (Table S1). A dose of rate of ∼13.872 e^-^/pix/sec was collected across 60 frames with a dose of ∼50 e^-^/Å^2^. To overcome problems of preferred orientation, the data was collected at the tilt of −20° (21). A total of 7289 micrographs (CiPer45: 1599; CiPer63: 2277; CiPer63PS: 1398; CiPer78: 2015) were collected.

### Electron microscopy data processing

The micrograph frames were aligned using MotionCor2 (22) within the Appion image processing wrapper (23). For all of the four CiPer datasets, the subsequent data processing was performed in cryoSPARC (24) (Figure S2). The CTF parameters were estimated using Patch CTF Estimation (25). Particles were auto-picked by blob-picker and extracted with default parameters. Several rounds of 2D Classification were performed to roughly exclude some 30S and 70S particles. Ab-initio Reconstruction and 3D Classification were performed to further classify the particles. Non-uniform Refinement was performed to get the final density maps.

The pairwise molecular weight difference was calculated among all the resolved density maps, and these differences were used as the metric for hierarchical cluster analysis, as described before (26). A 10.0 kDa threshold was chosen for identifying maps that were essentially similar, and the particles from such classes were combined by consensus refinement. Eventually, twenty-three distinct density maps were reconstructed from four datasets with global resolutions ranging from 4.7 Å to 8.4 Å. Figures of the density maps in this manuscript were prepared using UCSF ChimeraX (27).

### Quantitation of rRNA and protein occupancy in EM density maps

The crystal structure of *E. coli* 50S subunit (PDB: 4YBB) was segmented into 139 elements and binarized to serve as a reference surface (15,26). The density maps of iSAT time course intermediates were binarized with the threshold voxel level = 0.25. The relative values between the density maps and the reference surface maps were calculated, resulting in the occupancy values between 0 and 1 for each element. The occupancy values were hierarchically clustered using Euclidean distance with Ward linkage.

## RESULTS

### Circular permuted 23S rRNAs are competent for ribosome biogenesis in iSAT

As reported in the previous study, Δ7 *prrn* strain cells expressing circularly permuted rRNA exhibit only a slightly reduced growth compared to wild-type cells (7). According to the ribosome profiles reported previously, no ribosomal subunit intermediates were observed in the cells expressing CiPer45 and CiPer63 23S rRNAs. Therefore, it is challenging to purify and characterize the assembly intermediates directly from the cells. Our previous work has proved that the iSAT system could successfully mimic the ribosome biogenesis progress *in vivo* (5,6). The iSAT reaction can be halted at multiple time points, allowing the characterization of assembly intermediates at various stages through cryo-EM. By coupling with the transcription of rRNA, the assembly intermediates captured in the iSAT reaction may reflect the diversity of assembly intermediates compared to those that accumulate at steady-state. This enabled us to reconstruct the ribosome assembly landscape for the circularly permuted ribosomes.

Four plasmids encoding circularly permuted rRNAs were constructed based on the *E. coli rrnb* operon, as described in a previous report (7). In the CiPer45, CiPer63 and CiPer78 23S rRNAs, Helix 1 was “closed” by the insertion of a tetra loop. The processing stem (PS) at Helix 1 was removed except in CiPer63(+PS) 23S rRNA. To induce circular permutation, the Helix 1 and the processing stem were re-inserted at Helix 45, Helix 63 and Helix 78, respectively, in CiPer45, CiPer63 and CiPer63(+PS), and CiPer78. To ensure correct processing of the 23S rRNA at the newly created cleavage site, the selected helices are located at the outer surface of the 50S, which are expected to be accessible by RNase III.

The iSAT reactions were performed with the circularly permuted plasmids replacing the wild-type ones as the source of rRNA. As expected, the fluorescence signal of the four reactions with circularly permuted rRNAs have different levels of delay (Figure 1B), indicating that the assembly process was impaired, or that the translation of the assembled ribosomes was compromised, which needs to be furtherly determined by translation assays. The circularly permuted transcripts in iSAT reactions were extracted by Trizol, purified by columns, and analyzed by agarose gel electrophoresis. As shown in Figure 1C, the transcript purified from the iSAT reaction with the CiPer63(+PS) construction consists of two bands instead of a complete 23S rRNA, which is consistent with the previous report (7).

**Figure 1.**
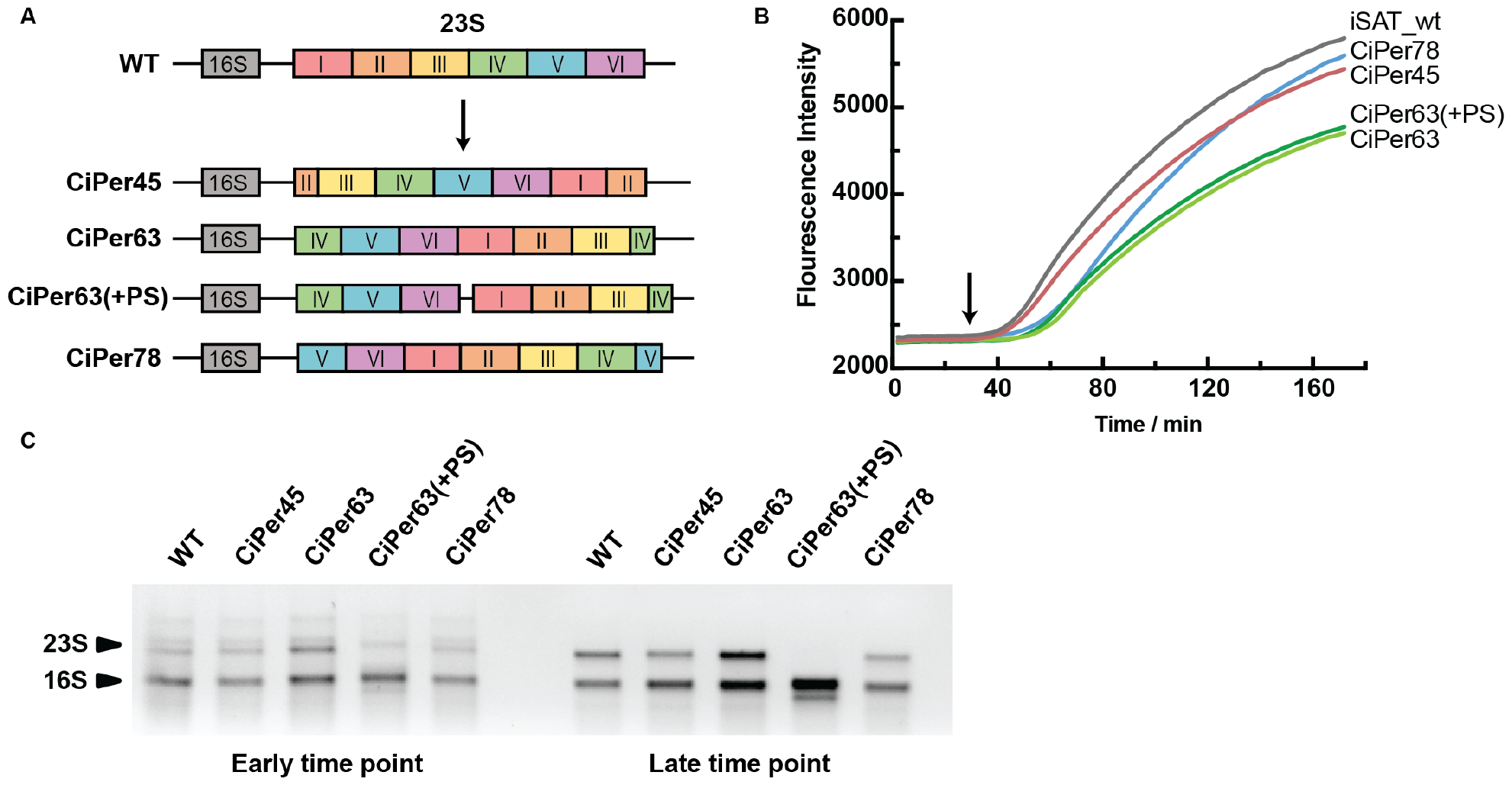
Circularly permuted rRNAs succeed in biogenesis in iSAT. **(A)** Schematics of the wild type and rearranged rRNA operons: CiPer45, CiPer63, CiPer63(+PS), and CiPer78. **(B)** The iSAT curve of circular permutated rRNA, reported by fluorescence signal of newly translated GFP. The reaction materials for the cryo-EM analysis were collected at the time point indicated by the arrow. **(C)** Agarose gel electrophoresis for the transcripts extracted from the iSAT reaction at the early time point (30 min) and late time point (120 min), respectively. The RNA product was purified by Direct-zol RNA Miniprep Kits of Zymo Research.

To capture of most immature intermediates, the iSAT reactions were quenched at an early time point as indicated by the arrow in Figure 1B. The iSAT reactions were analyzed by ultracentrifugation and sucrose gradient (Figure S1). The fractions containing 23S rRNA were concentrated and loaded onto grids. The cryo-EM data was collected on a Talos Arctica transmission electron microscope equipped with a K2 direct electron detector.

### Cryo-EM single particle analysis resolved structures of the circularly permuted LSU intermediates

The raw cryo-EM micrographs were processed using cryoSPARC in a similar manner to that previously described (24) (Figure S2). The particles were extracted from the filtered micrographs by Blob-Picker, and cleaned up by multiple rounds of 2D classification. After Ab-Initio Reconstruction, the particles that were successfully reconstructed into meaningful classes were homogeneously refined and then subclassified using 3D classification. The resolved classes were compared and combined by hierarchical analysis, as previously reported (26). Eventually, four ensembles of CiPer LSU intermediate structures were reconstructed from the four datasets, as shown in Figure 2, with resolutions range from 4.7 Å to 8.4 Å. Overall, the observed intermediates correspond well with previously observed intermediates, and consequently, the CiPer LSU intermediates are named and colored based on the superclasses defined in our previous work (15).

**Figure 2.**
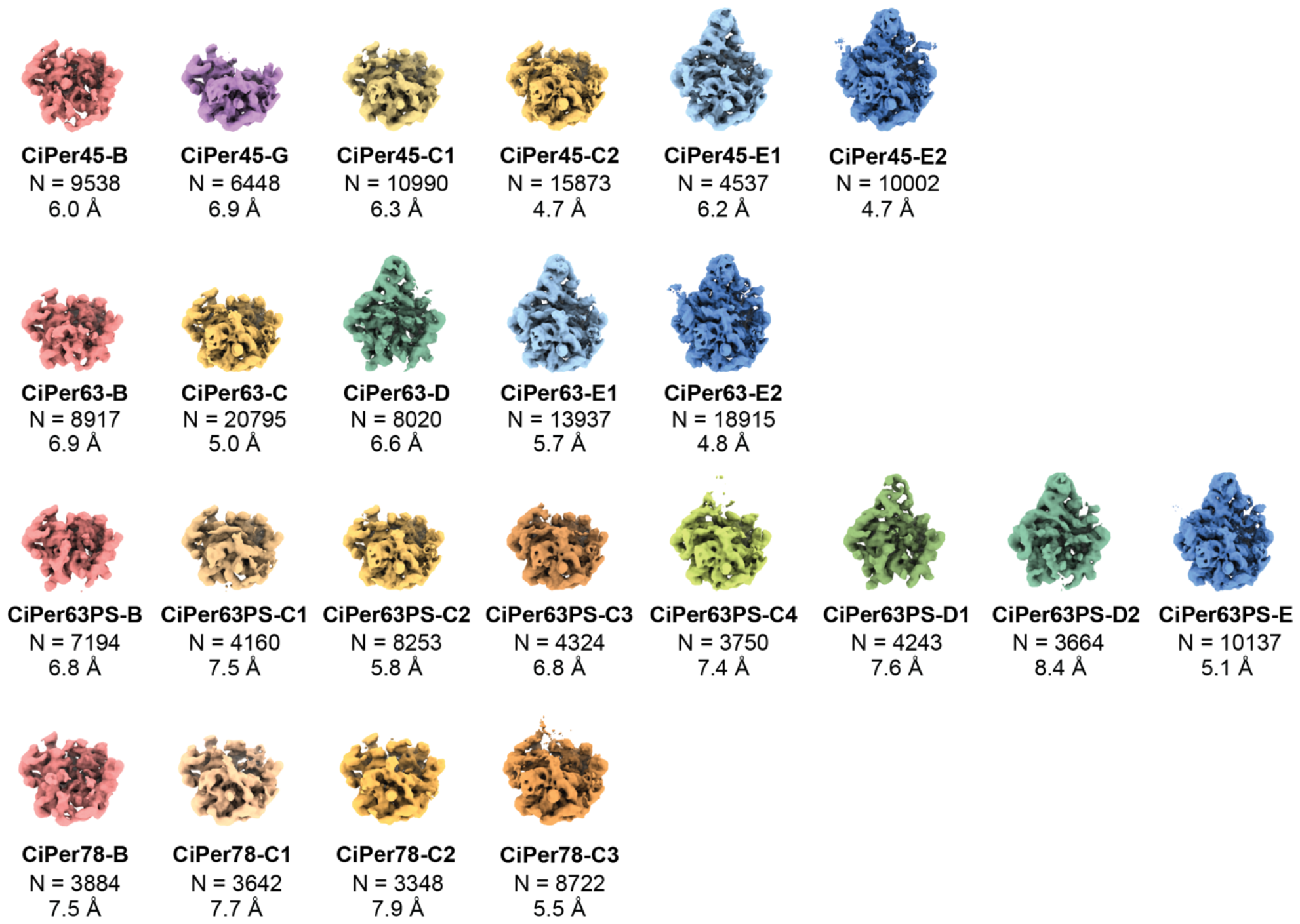
Density maps for circularly permuted LSU intermediates. Twenty-three 50S intermediates were obtained by heterogeneous subclassification from the iSAT reaction with circularly permutated rRNAs. The 50S precursors are named according to the major class they belong to, and are generally ordered from immature to mature. The number of particles contributing to each class is given along with the final resolution of the map.

To make the circular permutations in 23S rRNA, non-native sequences are inserted. Extra densities corresponding to these non-native regions are visible in the maps of the LSU intermediates. Figure 3 shows four B classes resolved from various CiPer datasets, each aligned with the atomic model of the mature 50S subunit. Extra electron density could be observed around Helix 45 in the CiPer45_B class (Figure 3A), likely attributable to the remnants of the processed inserted Helix 1 and processing stem. Similarly, an elongated segment could be observed around the native Helix 1 in the CiPer63(+PS)_B class, which is possibly the residue of the processing stem. These characteristics are exclusively present in their respective intermediates, serving to validate that these LSU intermediates are indeed composed of the circularly permuted 23S rRNAs.

**Figure 3.**
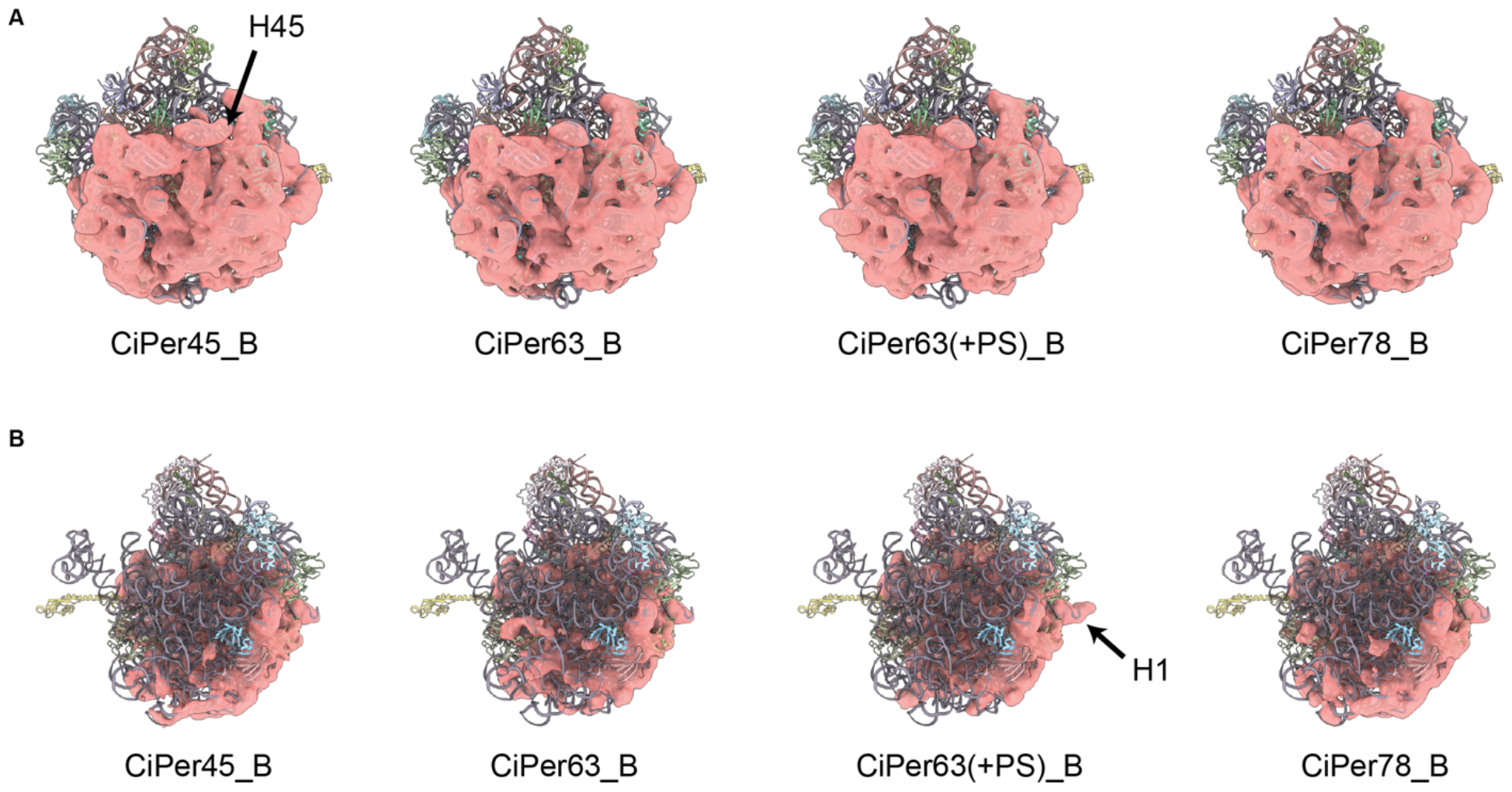
Circularly permuted LSU intermediates exhibit identified structural features. Four B classes in four CiPer datasets are shown in back **(A)** and side view **(B)** by the semi-transparent mask aligning with the atomic model of 50S subunit (PDB:4YBB). Arrows indicate the helices with extra density due to the circularly permuted rRNA constructs.

### Circularly permuted LSU intermediates exhibit specific structural features

The particle class distribution of the LSU intermediates in each CiPer dataset was analyzed (Figure S3). Intriguingly, several novel or rare classes were identified from CiPer45, CiPer63 and CiPer63(+PS) datasets. In iSAT time-course dataset, only about 2.5% particles were reconstructed into a G class intermediate for the early time points (5), while the proportion is 11.2% in CiPer45 dataset. The D class, which lacks density around around H63 in the base of the 50S, was not observed in the wildtype iSAT time-course intermediates, but was reported to constitute 20% of the particles in bL17-depletion datasets (15). In CiPer63 and CiPer63(+PS) dataset, 15.4% and 17.3% particles were identified as D classes, respectively.

The detailed structural composition of the reconstructed LSU intermediates was further analyzed by occupancy analysis, as shown in Figure 4A. As reported in previous work, by calculating the occupancy value of the resolved structures for each structural element, assembly blocks could be revealed, in which the structural elements behave similarly among the different structures as cooperative folding units (15).

**Figure 4.**
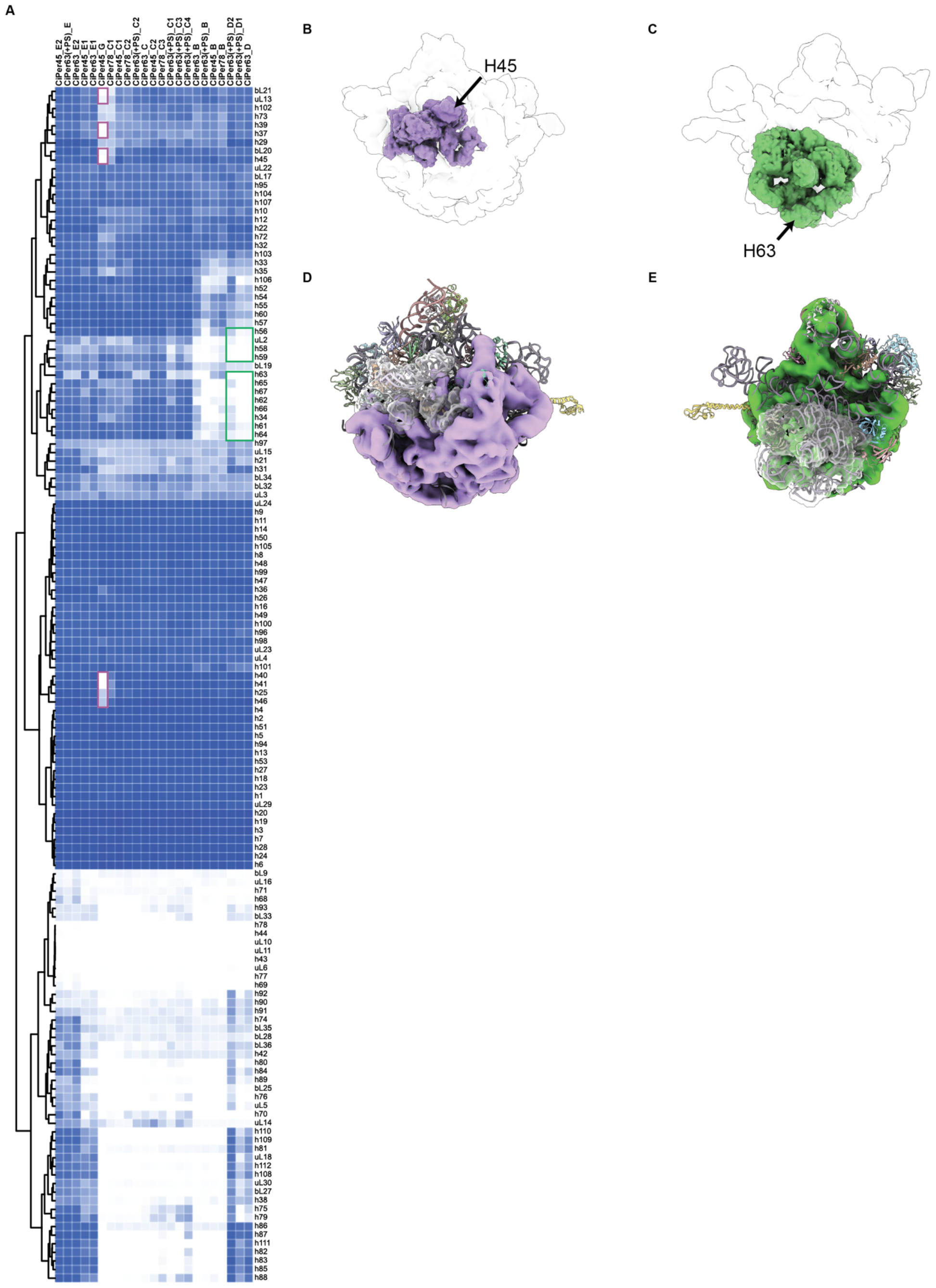
Certain structural elements are absent in circularly permuted LSU intermediates. **(A)** Occupancy of rRNA helices and ribosomal proteins. The values were used to analyze the correlation of the 139 assembly elements for each of the circularly permuted LSU intermediates density maps. The blue or white squares represent the presence or absence of the segments in the intermediates. The theoretical densities of the H45 Block **(B)** and H63 Block **(C)** are colored purple and green respectively in the 50S crystal structure (PDB: 4YBB). CiPer45_B **(D)** and CiPer63_D **(E)** classes are aligned with the 50S atomic model, with the H45 Block and H63 Block covered by a transparent mask, respectively.

The structural elements highlighted in purple rectangles in Figure 4A are exclusively absent in the CiPer45_G class. As depicted in Figure 4B, these elements are located in the vicinity of Helix 45, encompassing Helix 45. In the construction of CiPer45 23S rRNA, the 5’ and 3’ ends of Helix 45 were separated to the two ends of the 23S rRNA, which means Helix 45 can only form after the complete transcription of the entire 23S rRNA. Consequently, the absence of the purple elements referred to as the “H45 Block” (Figure 4D) is likely caused by the perturbation of Helix 45, suggesting an interdependency between these purple elements and Helix 45. Similarly, the absence of H63 Block around Helix 63 is observed in CiPer63_D, CiPer63(+PS)_D1 and CiPer63(+PS)_D2 classes (Figure 4C and 4E).

The Helix 78 is located in the L1 stalk, which is flexible and rarely be resolved by cryo-EM single particle analysis, making it hard to observe the folding of Helix 78 in CiPer78 LSU intermediates. But an interesting feature of the CiPer78 LSU intermediates is that they exclusively contain B and C classes, with no presence of E classes (Figure S3). This observation suggests that these CiPer78 LSU precursors may encounter difficulties in docking of the central protuberance. Overall, delayed transcription of specific helices in circularly permuted 23S rRNAs gives rise to locally distinct intermediates. These intermediates further elucidate relationships among the structural elements associated with or reliant upon the perturbed helices.

### Circularly permuted 23S rRNAs follow the native co-transcriptional docking hierarchy with limited perturbation

Based on the structural composition of the circularly permuted LSU intermediates, the resolved structures are arranged in a sequence to illustrate the assembly pathways involved in the biogenesis of the circularly permuted LSUs, as depicted in Figure 5. Arrows are used to denote the minimal conformational changes between the intermediates, from immature to mature states. In comparison with our previous report (5), most circularly permuted LSU intermediates follow the original B-C-E pathways. Not all the intermediates observed in the wild-type iSAT time-course dataset are present in the CiPer datasets, potentially due to the lower number of particles in the CiPer datasets. In addition, the emerging assembly challenges existing in the circularly permutants may also result in the class repopulation.

**Figure 5.**
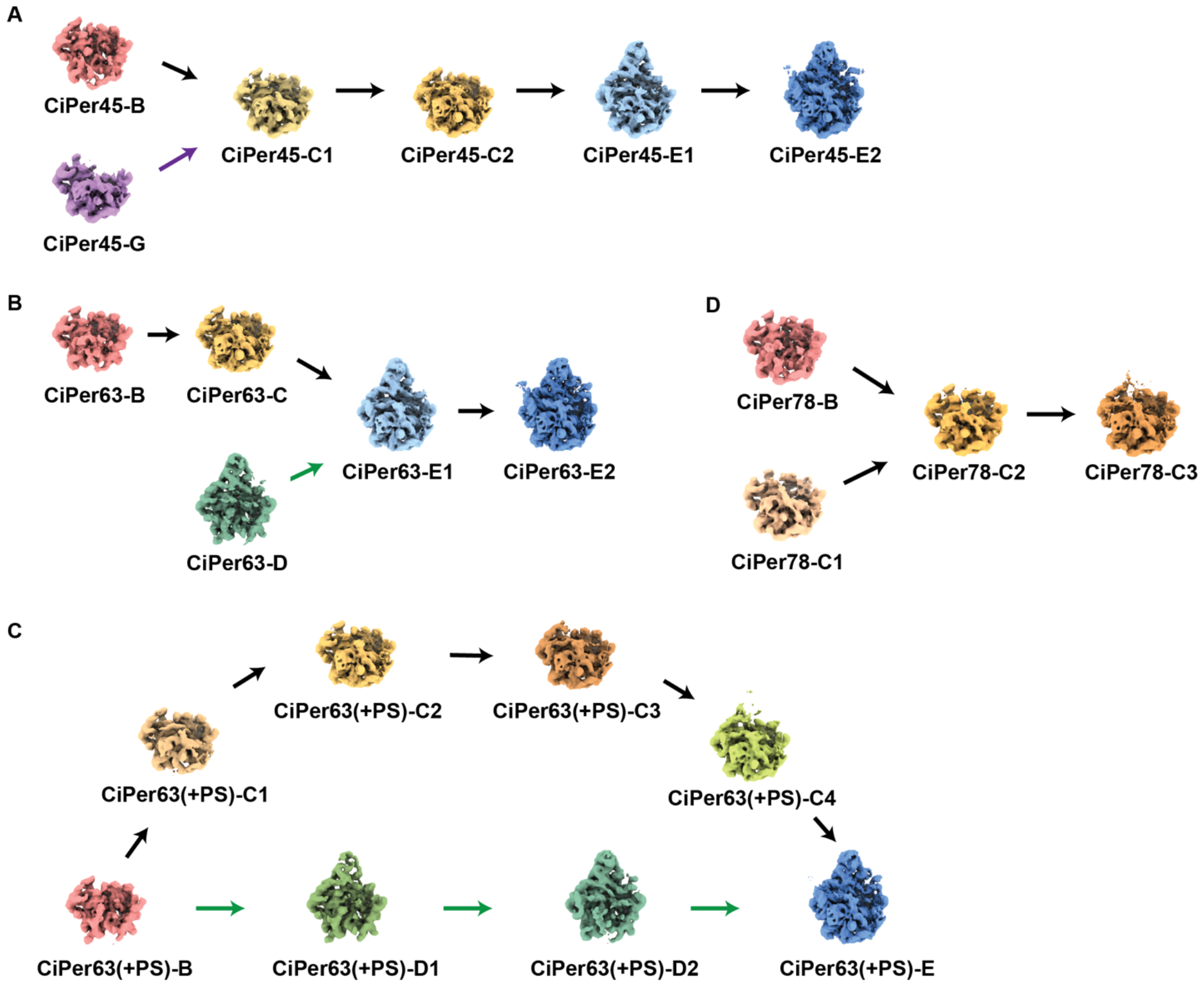
Organization of circularly permuted LSU intermediates into assembly pathways. The reconstructed 50S intermediates from CiPer45 **(A)**, CiPer63 **(B)**, CiPer63(+PS) **(C)** and CiPer78 **(D)** are organized according to the composition analysis. The arrows indicate the minimal folding steps required to connect the set of intermediates. The purple and green arrows indicate the parallel assembly pathways caused by the perturbation of the split helices in CiPer45 and CiPer63 rRNA constructs.

Parallel pathways resulting from the disruption of split helices are evident in the biogenesis of the circular permutants. While some circularly permuted LSU intermediates lack the affected blocks, the intermediates that were previously observed in wild-type datasets do exist, forming the parallel pathways. This indicates that the disruption of split helices may not necessarily affect every circularly permuted 23S rRNA molecule. Taking the CiPer63(+PS) dataset as an example, when the central protuberance docks before the formation of the base portion, it is observed as a D class. Similarly, the C classes are resolved when the docking of the base part occurs before the formation of the central protuberance. Since both C and D classes are observed in the CiPer63 and CiPer63(+PS) datasets, the docking challenges encountered by the central protuberance and the base part do not significantly differ.

In conclusion, while several novel circularly permuted LSU intermediates are resolved in the CiPer datasets, the assembly pathways observed for circularly permuted 23S rRNAs during biogenesis generally align with those observed in the wild-type dataset. This suggests that the docking hierarchy plays a predominant role as the main driving force, even when the docking hierarchy conflicts with the 5’-3’ co-transcriptional directionality (Figure 6).

**Figure 6.**
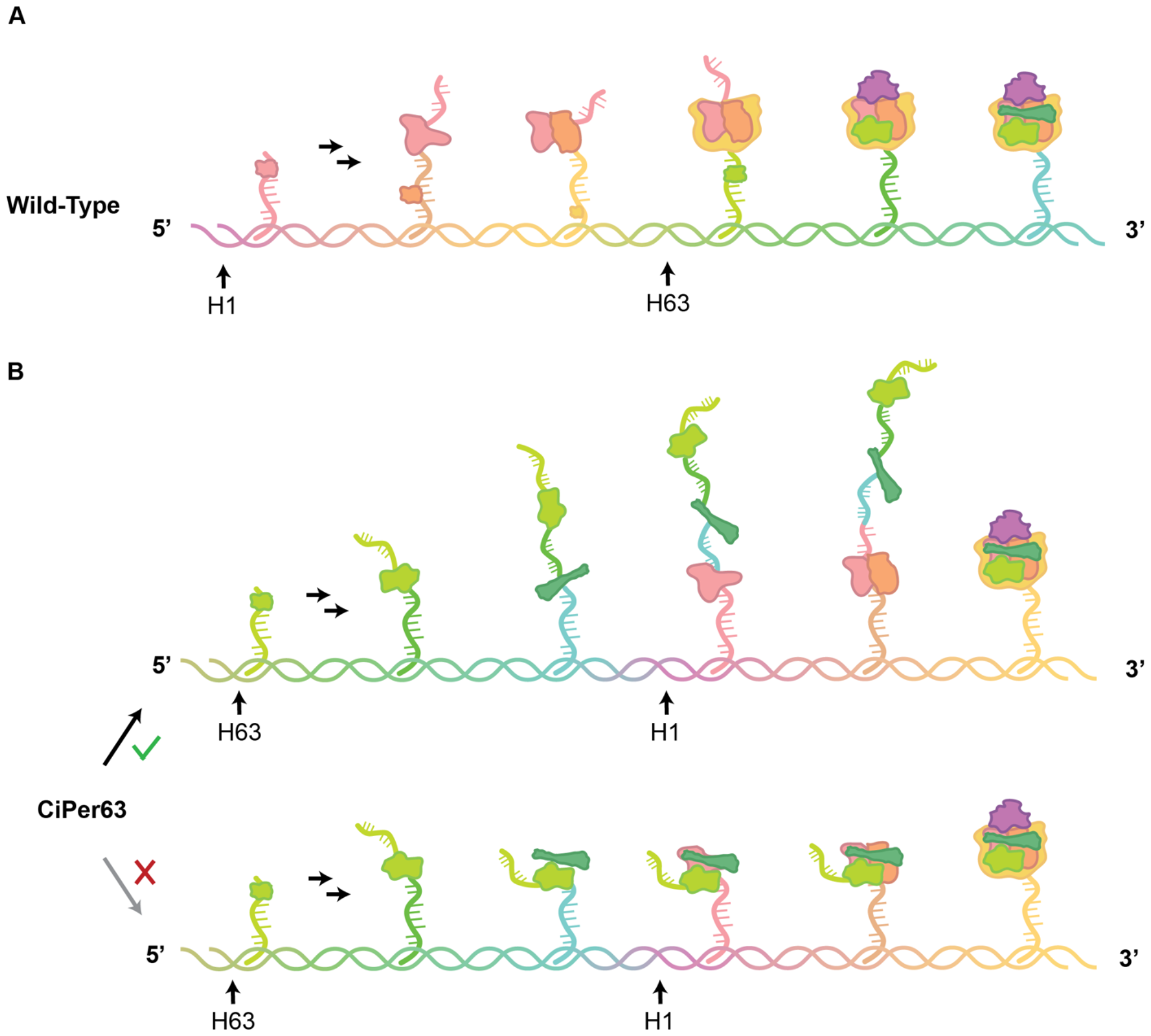
The docking hierarchy drives rRNA folding predominantly. **(A)** Simplified schematic diagram illustrating the assembly process of wild type 23S rRNA. The assembly intermediates at various stages are listed alongside the transcription directionality from the 5’ end to the 3’ end. Consistent color schemes are applied to DNA and RNA strands. Irregular polygons depict the docked assembly blocks composed of folded rRNA and ribosomal proteins. With the state-of-the-art single-particle cryo-EM analysis, only the docked portions are sufficiently rigid to be resolved. **(B)** Assembly of circularly permuted 23S rRNA with CiPer63 presented as an example, showcasing two mechanisms governed by different logics. In the top mechanism, validated in this study, rRNA can fold and assemble into modules concurrently with transcription but docks hierarchically, beginning with the pink module. In the bottom mechanism, rRNA folds and docks in tandem with transcription from the 5’ to 3’ end, which could begin with any module.

## DISCUSSION

According to previous studies, the folding of bacterial rRNA has been proposed to occur co-transcriptionally from the 5’ to 3’ end (1-3). Further, our laboratory has previously reported a modular assembly mechanism (5,6,15), wherein rRNA helices fold cooperatively with ribosomal proteins and dock as blocks onto assembly precursors in a hierarchical manner. For wild-type rRNA constructs, this docking hierarchy generally aligns with the directionality of transcription, especially for the early assembly stages (4-6) (Figure 6A). However, in circularly permuted rRNA constructs, the transcription order is rearranged, providing a context to investigate the predominant intrinsic forces governing rRNA folding (Figure 6B).

If rRNA undergoes folding and assembly in a strict co-transcriptional order, it is anticipated that alternative assembly cores and rearranged assembly pathways would be observed (Figure 6B). For instance, in the case of the CiPer63 23S rRNA construct, a precursor composed solely of Domain III or III+IV of the 23S rRNA is expected to form and be resolved within the context of co-transcriptional folding. Consequently, the rRNAs will undergo biogenesis through entirely different assembly pathways compared to those observed previously. In this study, we reconstituted ribosomes using circularly permuted rRNAs in near-physiological iSAT reactions and employed cryo-EM single particle analysis to characterize the LSU assembly intermediates present in the iSAT reaction. Remarkably, no intermediates with alternative assembly cores were identified. All circularly permuted LSU intermediates were found to adhere to the known docking hierarchy, notwithstanding some unique classes resulting from the disruption of split helices. However, concurrently, an alternative possibility exists, wherein the docking of the assembly blocks is faster than folding. Subsequently, upon folding, the docking process transpires rapidly, thereby obscuring the intermediary steps.

We must acknowledge that this finding could be constrained by current technological limitations. Resolving the alternative assembly core or precursors may prove challenging for several reasons. Firstly, the size of the alternative assembly core could present a limitation. Based on our previous studies, the smallest core identified by cryo-EM contains 600 nucleotides. Therefore, cores potentially existing below this threshold might be too small to be aligned and resolved during heterogeneous reconstruction. Moreover, given that cryo-EM single-particle analysis can only resolve densities that are not highly flexible, we cannot conclusively determine whether certain rRNA helices have formed but not yet docked into higher-order structures. It seems plausible that the circularly permuted 23S rRNA still undergoes co-transcriptional local folding into secondary structures, albeit adhering to the original docking hierarchy for assembly into tertiary structures. Nonetheless, we have made diligent efforts to thoroughly explore the database as comprehensively as possible. We abstained from using any template references during particle picking and consequently reconstructions, thus ensuring that the strategies employed during data processing have been implemented with minimal bias. As recently reviewed (28,29), alongside single-particle cryo-EM analysis, emerging biophysical approaches such as single-molecule fluorescence microscopy and RNA structure probing have been developed to further investigate the assembly mechanisms of RNA-protein complexes. A more comprehensive understanding could be attained through the integration of additional methods in future studies.

The predominance of the docking hierarchy in rRNA folding, rather than the 5’-3’ co-transcriptional directionality, is reasonable for several reasons. Firstly, it is imperative to ensure that rRNA forms correct intermediate conformations, which are supposed to be specifically recognized by ribosomal proteins (30-32) or assembly factors (29,33-36). Secondly, the 5’ to 3’ directionality is not always rigorously adhered to in the whole biogenesis process of the wild type 23S rRNA. For instance, to ensure the translation fidelity, the peptidyl transferase center (PTC) serves as the final assembly block that docks into the functional conformation. This observation also indicated the dominant role of the docking hierarchy in ribosomal LSU assembly. Additionally, in extreme cases such as *Chlamydomonas reinhardtii* (37), rRNA is arranged into a scrambled genome. Given the typically high conservation of ribosomal structures among various strains (38-40), the robust docking hierarchy of rRNA offers theoretical plausibility for the successful biogenesis of such a scrambled rRNA genome.

Among the four circularly permuted 23S rRNAs, the CiPer63(+PS) was designed to be processed into two rRNA fragments by retaining the processing stem instead of replacing it by a tetraloop. In both previous *in vivo* and our *in vitro* studies, no severe defects were observed with the biogenesis of CiPer63(+PS) rRNA. Unlike truly fragmented rRNA constructs, processing of CiPer63(+PS) 23S rRNA could only occur after the elongated CiPer63 helix, which serves as the recognition sequence of RNase III, has formed. Consequently, although we have determined that the CiPer63(+PS) 23S rRNA is indeed processed into two fragments in the iSAT reaction, no significant perturbation actually exists during its folding. Thus, no significant differences were observed in its biogenesis compared to CiPer63 23S rRNA.

According to the previous report, cells containing CiPer78 23S rRNAs displayed severe growth defects (7), whereas its biogenesis rate is similar to that of CiPer63 and CiPer63(+PS) in the iSAT reaction. This difference may be attributed to the fact that the iSAT reaction, despite operating under near-physiological conditions, may not fully reflect stresses that occur *in vivo*. In the CiPer78 23S rRNA construct, the PTC segment is transcribed first, while it is the last part to be docked according to the assembly hierarchy. During the unusual prolonged “waiting period”, the 5’ of the transcript could be recognized and partially degraded by quality control factors in the cell due to its abnormal solvent exposure.

In the original report of circularly permuted rRNAs, the DEAD box RNA helicase *deaD* (*csdA*) was shown to play a crucial role in the biogenesis of circularly permuted ribosomes at 37 °C (7). In the wild type cells, DeaD was found to be a cold shock protein, which is indispensable for bacterial growth only at low temperatures, whereas its absence at 37 °C had no effect (41-43). Our previous study has shown an ensemble of LSU intermediates generated from the Δ*deaD* cells (6). The resolved LSU intermediates, trapped due to the absence of DeaD, represent states at a very early assembly stage, suggesting that DeaD participates in biogenesis from the outset. In the case of circularly permuted rRNAs, the primary structure of the rRNA operon is dramatically changed. The altered rRNA sequence context in circularly permuted rRNA could pose challenges for the correct folding of the early assembly core, thereby emphasizing the crucial role of DeaD.

In conclusion, our study marks the first report of intermediate structures of the ribosome large subunit containing circularly permuted 23S rRNAs, shedding light on their folding mechanisms. These novel findings underscore the dominant significance of the docking hierarchy among assembly blocks, taking precedence over the 5’-3’ co-transcriptional directionality. Our research contributes to a deeper understanding of the inherent logic of the rRNA folding and the intricate molecular mechanisms governing ribosome biogenesis.

## DATA AVAILABILITY

The density maps have been deposited in the EMDB with the following accession numbers: CiPer45-B: EMD-43844, CiPer45-C1: EMD-43845, CiPer45-C2: EMD-43846, CiPer45-E2: EMD-43847, CiPer45-G: EMD-43848, CiPer45-E1: EMD-43849, CiPer63-C: EMD-43850, CiPer63-E2: EMD-43851, CiPer63-E1: EMD-43852, CiPer63-D: EMD-43853, CiPer63-B: EMD-43854, CiPer63(+PS)-E: EMD-43856, CiPer63(+PS)-C2: EMD-43857, CiPer63(+PS)-B: EMD-43858, CiPer63(+PS)-C3: EMD-43859, CiPer63(+PS)-D1: EMD-43860, CiPer63(+PS)-C1: EMD-43861, CiPer63(+PS)-C4: EMD-43862, CiPer63(+PS)-D2: EMD-43863, CiPer78-B: EMD-43864, CiPer78-C1: EMD-43865, CiPer78-C2: EMD-43866, CiPer78-C3: EMD-43867.

