## Supplementary Materials for "Assembly of the Bacterial Ribosome with Circularly Permuted rRNA"

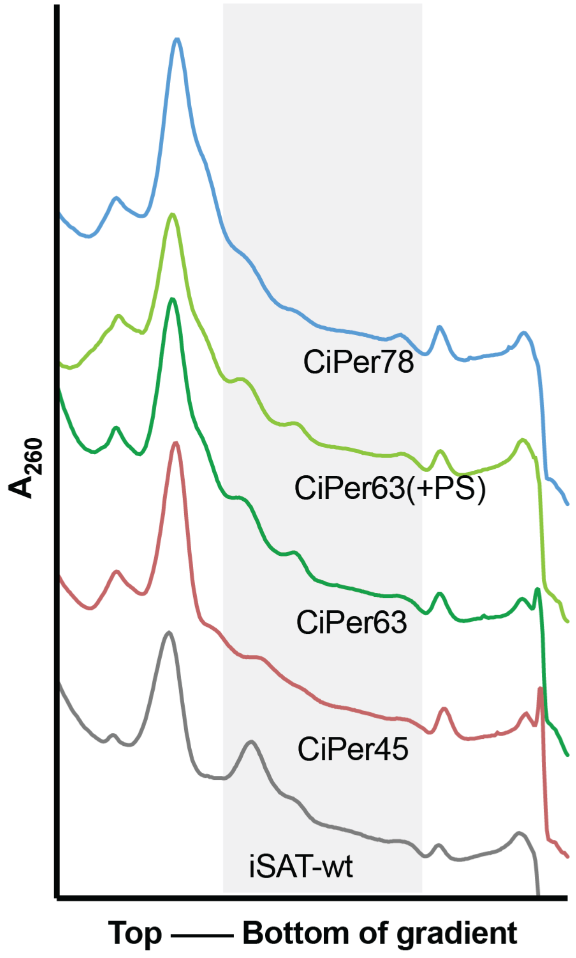


**Figure S1. Ribosome profile of iSAT reactions with circularly permuted 23S rRNAs.**

Four circularly permuted iSAT reactions were quenched on ice at early time point described in Figure 1. The ribosome profiles of the iSAT reactions were analyzed using sucrose density gradient ultracentrifugation. The fractions highlighted by the grey shadow were collected and concentrated for the cryo-EM data collection.

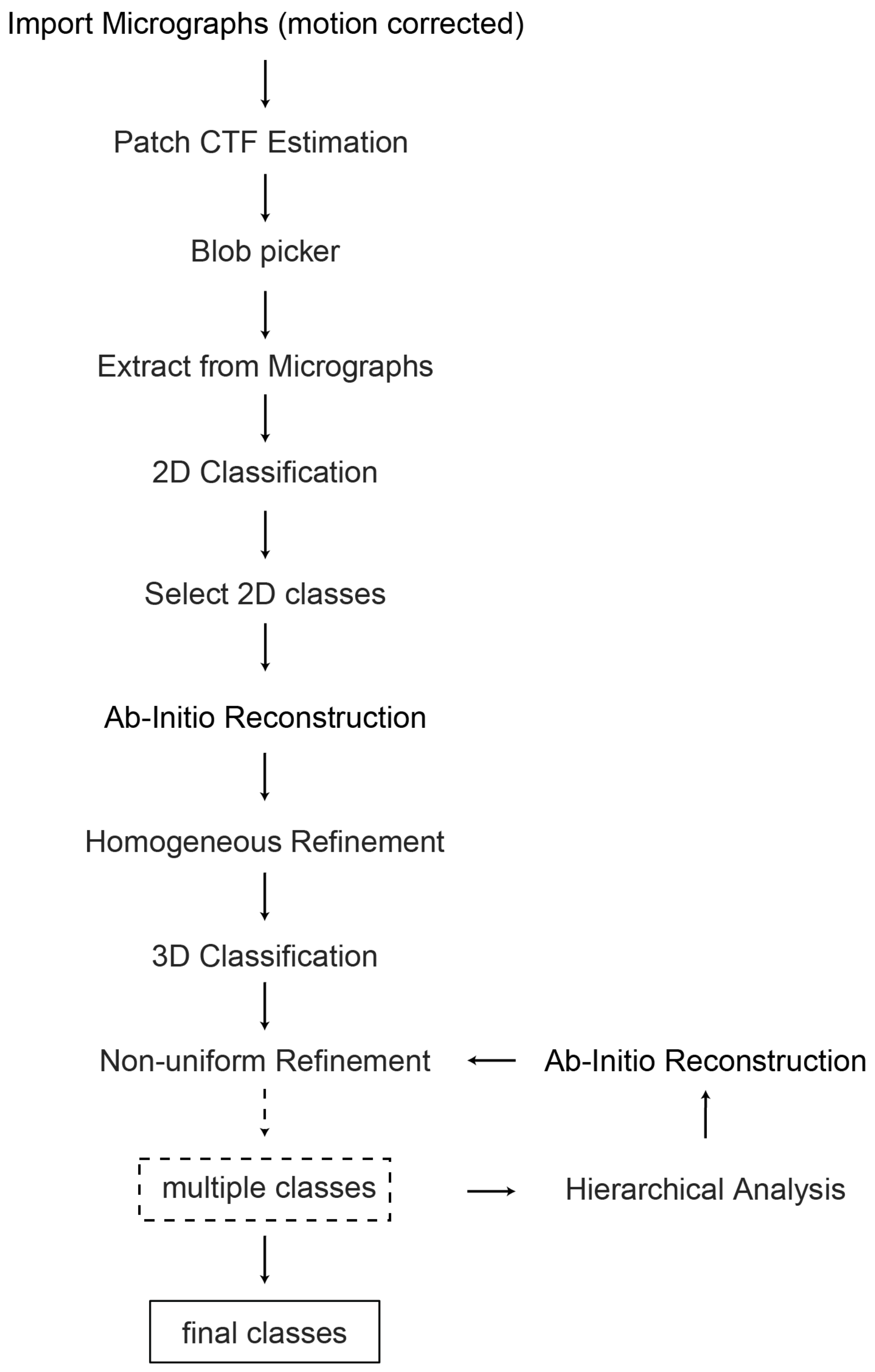


**Figure S2. Workflow of cryo-EM single-particle analysis by cryoSPARC.**

The circularly permuted LSU intermediates are processed by the workflow of cryo-EM single-particle analysis indicated here. More details are described in Methods.

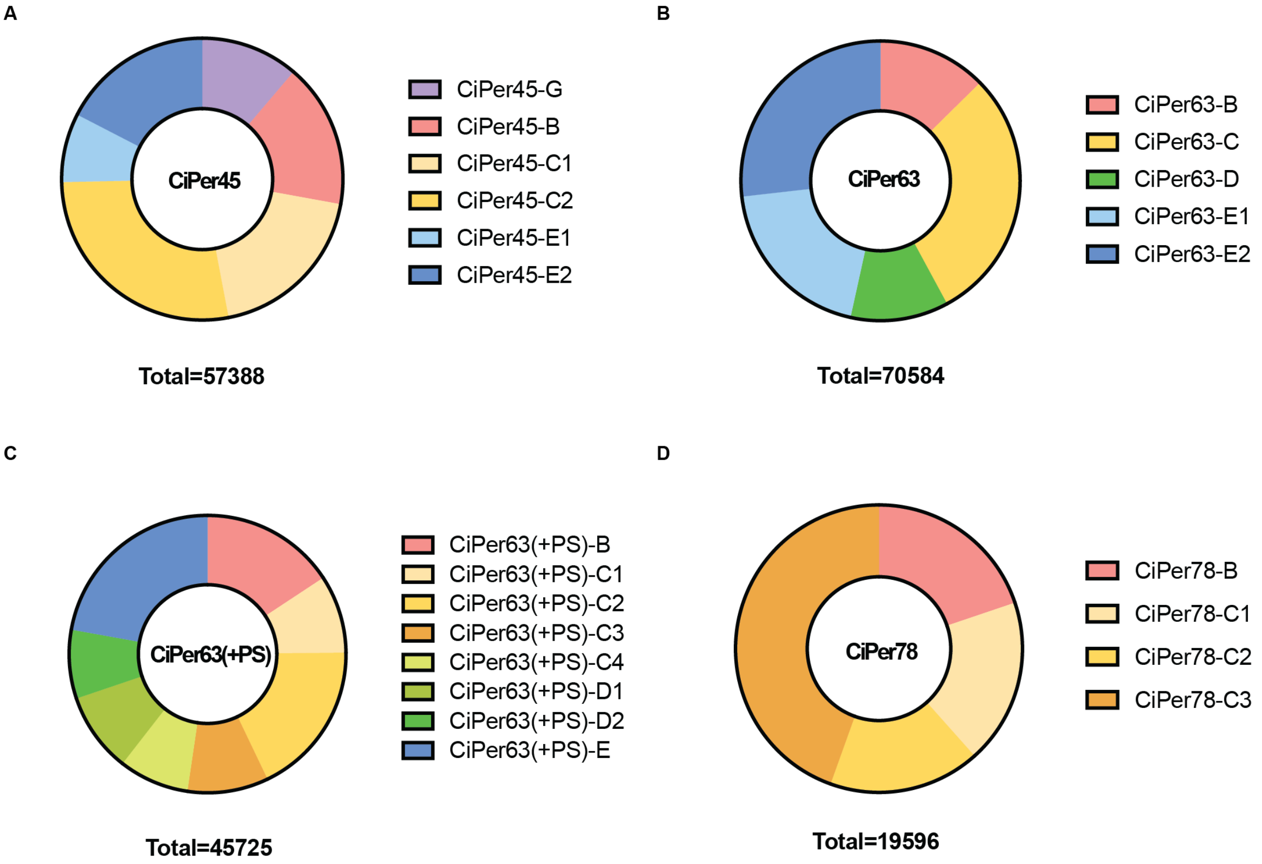


**Figure S3. Particle class distribution for circularly permuted LSU intermediates.**

The population distributions for the CiPer datasets were calculated and plotted in pie chart. The color code is consistent with that in Figure 2.

**Table S1. CryoEM data collection and image processing parameters**

| CryoEM data | | | | | |
|---|---|---|---|---|---|
| **Data acquisition** | | **CiPer45** | **CiPer63** | **CiPer63(+PS)** | **CiPer78** |
| Microscope | | FEI TALOS ARCTICA | | | |
| Camera | | Gatan K2 Summit | | | |
| Magnification | | 36000 | | | |
| Stage Tilt (°) | | -20 | | | |
| Voltage (kV) | | 200 | | | |
| Micrographs | | 1599 | 2277 | 1398 | 2015 |
| Exposure time (ms) | | 4800 | | | 5000 |
| Number of frames | | 24 | | | 25 |
| Dose rate ($e^-$ /pixel /s) | | 13.872 | | | 13.062 |
| Total Exposure ($e^-$ /Å$^2$) | | 50 | | | |
| Defocus range (µm) | | -1.0 to -2.5 | | | |
| Pixel size (Å /pixel) | | 1.15 | | | |
| Box size (pixel) | | 366 | | | |
| Symmetry imposed | | C1 | | | |
| Number of particles | Extracted | 487,886 | 686,910 | 413,898 | 338,996 |
| | After 2D Classification | 263,718 | 228,908 | 148,997 | 47,947 |
| | After Ab-Initio Reconstruction | 60,382 | 81,709 | 50,952 | 19,596 |
| | After 3D Classification | 57,388 | 70,584 | 45,725 | 19,596 |

| FSC threshold | 0.143 | | | |
|---|---|---|---|---|
| Resolution range (Å) | 4.8-6.8 | 4.9-7.0 | 5.3-8.7 | 5.6-8.7 |
